# The temporal and perceptual characteristics of emotion-induced blindness

**DOI:** 10.1101/2025.05.15.653853

**Authors:** Michaela E. Alarie, Grace H. Yang, Lila R. Quinn, Tiffany Lin, Wael F. Asaad

## Abstract

Attentional capture by emotionally salient stimuli is adaptive, permitting identification of possible threats; however, an excessive bias towards emotional stimuli can interrupt goal-directed behavior. This is especially relevant in psychiatric disease, where severe emotional distress can interfere with daily function. As such, understanding the mechanisms by which emotional stimuli compete for attentional resources is a critical area of investigation. Previous studies using rapid serial visual presentation (RSVP) paradigms observe that emotional distractors disrupt the detection of subsequent stimuli, referred to as emotion-induced blindness (EIB). Our study expands upon this work, characterizing how temporal and perceptual factors shape the emergence and intensity of EIB. Contrary to previous assumptions regarding temporal dynamics of EIB, we found that effects of emotional distractors persisted across prolonged image presentation durations. Further, we investigated the extent to which the depth of distractor processing influences EIB using a distractor recall task. While recall was predictive of EIB magnitude, a significant effect of emotional distractors on target detection was nonetheless present even without conscious recall of the distractor. These findings demonstrate the robustness of the EIB effect in RSVP in the context of temporal and perceptual manipulations.

## Introduction

Attending to stimuli in a rapidly changing environment has survival value. In particular, recognizing emotionally salient stimuli that present a potential threat is necessary to generate behaviorally relevant actions. However, emotional stimuli might also impair goal-directed behaviors by capturing limited attentional resources. This is relevant in the context of psychiatric disorders, which often involve an excessive bias towards emotionally-salient stimuli, leading to deficits in daily function (Bredemeier et al., 2011; Olatunji, 2021; Tonta et al., 2023; Waszczuk et al., 2015). As such, understanding the mechanisms by which emotional stimuli capture attention is a critical area of investigation (McHugo et al., 2013; Wang et al., 2012).

Previous work has demonstrated that emotionally salient distractors impair perception of subsequent stimuli, formally referred to as emotion induced blindness (EIB) (Goodhew & Edwards, 2022; McHugo et al., 2013). Many EIB studies have harnessed rapid serial visual presentation (RSVP) tasks (Kennedy & Most, 2011; Kennedy & Most, 2015; Most et al., 2005; Most & Jungé, 2008). In one typical EIB variant of RSVP, participants search amongst a stream of 17 landscape/architectural photos for an image that is rotated 90 degrees. At the end of each trial, participants are instructed to report the orientation of the rotated image in the stream. Depending on the trial, a task-irrelevant negative or neutral distractor image precedes this rotated target. Affectively-negative distractors, in particular, impair the ability to report the rotated target.

Additional work has delineated some of the factors that most strongly modulate EIB. For example, manipulations of the sequential proximity between emotional distractors and target stimuli attenuate the EIB (Most et al., 2005). Less proximity between these two stimuli, that is, greater lag between the emotional distractor and the target, attenuate EIB. The effect of emotionally salient distractors persisted when emotional distractors were shown up to 4 images preceding the target (about 400–600 ms earlier) (Ciesielski et al., 2010; Most & Jungé, 2008).

While various behavioral mechanisms of EIB have been studied, open questions remain. For example, the majority of EIB studies employ a constant image presentation duration (e.g.,100 ms per image), utilizing lag manipulations to probe temporal dynamics. However, this approach conflates temporal distance with the number of intervening stimuli, limiting insights into the impact of stimulus duration alone. Another open question concerns the depth of processing required for emotional distractors to elicit EIB. While it has been rigorously demonstrated that emotional distractors impair target recall, it is unclear whether participants must be consciously aware of the distractor’s content for this interference to occur. A more complete account of the behavioral mechanisms underlying EIB requires disentangling these contributing factors to determine the specific task demands and perceptual conditions under which EIB emerges.

Our present study aims to further characterize the conditions required for affectively charged stimuli to impair visual attention. In experiment 1, we validated an adapted RSVP paradigm for investigating EIB, ensuring results aligned with prior reports. In experiment 2 we investigated the role of temporal dynamics in modulating EIB severity through manipulations of image presentation duration. Finally, in experiment 3 we assessed the depth of processing of emotionally salient distractors required to trigger EIB. Together, findings from this study refine our understanding of the behavioral mechanisms underlying EIB, offering further insight into how emotionally salient stimuli capture visual attention.

## Materials & Methods

### Participants

A total of 87 participants were recruited from Brown University and the broader Rhode Island communities. Participants provided written informed consent prior to beginning the study and had normal or corrected-to-normal vision. The study was approved by the Institutional Review Board at Rhode Island Hospital, and all experiments were performed in accordance with the guidelines and regulations set by this institution. Participants were excluded if they had a history of moderate to severe psychiatric illness.

All subjects first performed the titration phase of the experiment. Following titration, subjects performed either experiment 1, 2, or 3. A total of 22 (5 male, 17 female, mean age = 21.8 years, range 18–43 years), 41 (11 male, 30 female, mean age = 19.3 years, range 18–21 years), and 30 (12 male, 18 female, mean age = 21.6 years, range 18–31 years) participated in experiments 1-3, respectively. N=2 completed experiments 1 and 3, N=9 completed experiments 2 and 3, and N=3 completed all 3 experiments. Participants who completed multiple experiments did so on separate days, repeating titration before the start of each experiment. Sample sizes for each experiment were comparable to previous studies of EIB (Kennedy & Most, 2015; Singh & Sunny, 2017).

### Experimental Setup and General Procedure

Stimuli were presented on a 27-inch LED monitor (resolution 1920 × 1080 pixels, refresh rate 60 Hz) using the NIMH MonkeyLogic toolbox for MATLAB (Asaad et al., 2013; Hwang et al., 2019). Subjects were seated approximately 57 cm from the monitor. Stream and target images were drawn from the International Affective Picture System (IAPS) and eBird databases, respectively (Bradley & Lang, 2017; eBird, 2021; Sullivan, 2009). All images were 300 × 300 pixels in size and presented at the center of the screen against a dark background. All experiments were performed in a well-lit room. Detailed information on the procedure for each experiment is included below, with a summary in Table 1.

**Table 1.** Experiment Procedure and Metadata.

|  | Image Duration (ms) | Distractor Lag from Target | Distractor Types | Trials per Condition | Recall phase | N |
| --- | --- | --- | --- | --- | --- | --- |
| Experiment 1 | 100 | 1, 2, 8 | Neutral, negative, scrambled negative | 16 | N | 22 |
| Experiment 2 | 33.33, 50, 100, 150, 200, 300 | 1 | Neutral, negative | 30 | N | 41 |
| Experiment 3 | 83.33, 100, 200, 300* (n=18) | 1 | Neutral, negative | 40 | Y | 30* |

All trials consisted of an RSVP stream of 17 temporally-contiguous images (0 ms inter-image-interval; Fig. 1A). Trials were initiated by pressing a button, at which time a fixation spot appeared for 1000 ms, followed by the RSVP image stream, leading immediately into the choice phase. Each trial contained one target: a trial-unique image of a bird selected at random from the database of bird images. Participants were instructed to search for this bird image within the stream in order to correctly identify it during the choice phase. Subjects used a joystick-controlled cursor to select the target bird image from among six bird image options (Fig. 1B); chance performance was 16.67%. The chosen image was outlined in green if correct and red if incorrect. If the incorrect option was chosen, the correct image was concurrently outlined in green. Experiments varied by probe distractor type (Fig. 1C), image presentation duration, and the inclusion of a recall phase. Probe distractors were salient images presented before the target and belonged to one of three categories: negative, scrambled negative, or neutral. There were 22 negative, 22 scrambled negative, and 92 neutral distractor images. Probe distractors were selected randomly and replaced only once all had been used.

**Figure 1.**
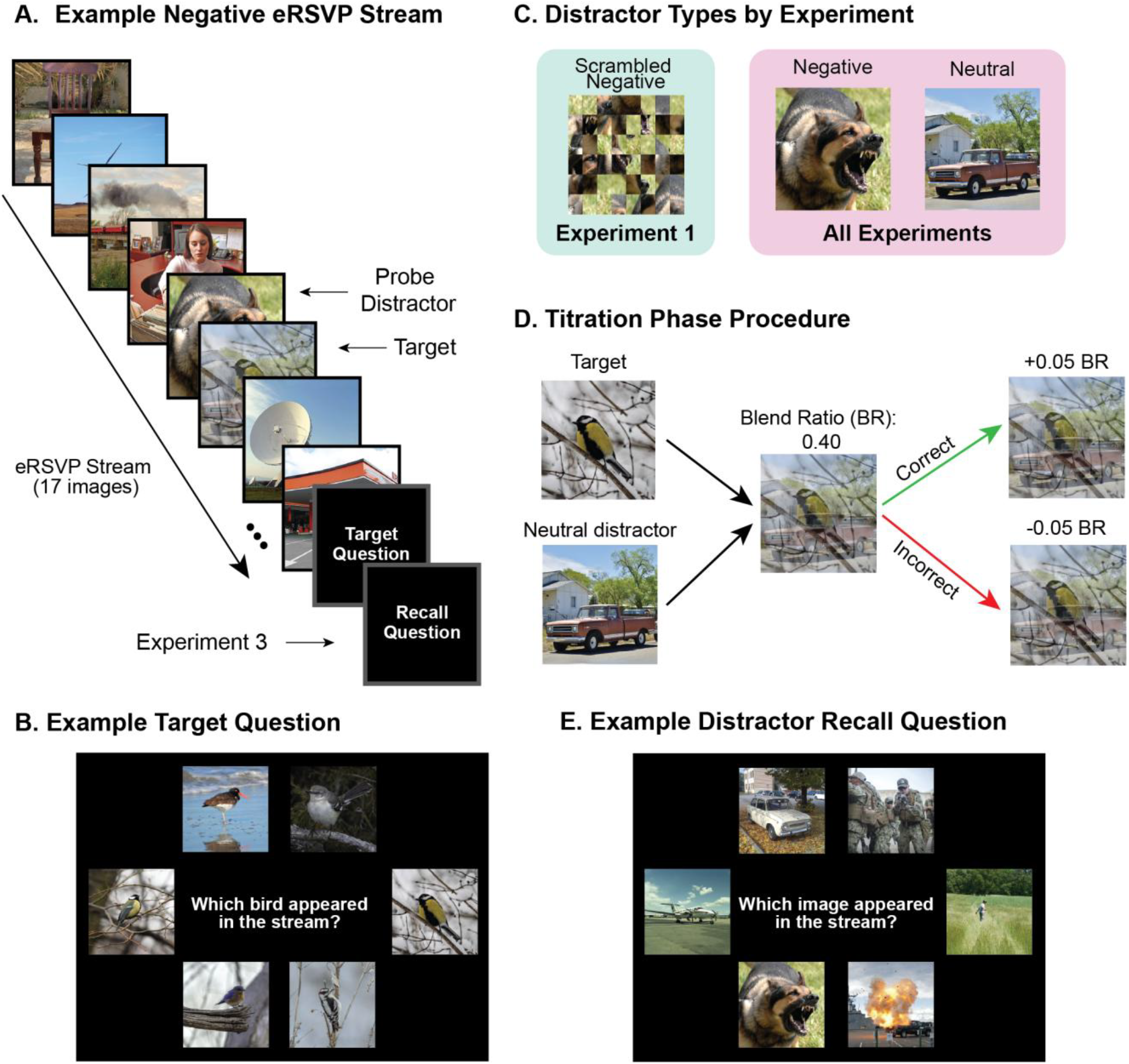
***A)***Example of emotional rapid serial visual presentation stream. Participants are instructed to scan the stream for the target, a single image of a bird. Participants are then asked which target (all experiments) and image (experiment 3) appeared in the image stream. ***B)***Example target question following the stream, referred to as the 5000 ms choice phase. ***C)***Example distractors across experiments. ***D)***Titration example for modulating task difficulty. All participants begin at a 0.25 blend ratio (BR) for the target image. Every successful trial resulted in a 0.05 increase in BR (and vice versa for every failed trial). ***E)***Example recall question following the target question, referred to as the 5000 ms recall phase. **A-E)** All images in this figure represent examples and are not from the eBird or IAPS databases. Images are public domain, sourced from Wikimedia Commons and Openverse.

### Titration Phase

Task difficulty was adjusted by blending the image of the target bird with a neutral distractor image to varying degrees, calibrated for each participant during the titration phase. Titration consisted of 75 trials, with each image presented for a duration of 100 ms. Participant-specific task difficulty was adjusted by blending the target image with a neutral distractor, referred to as blend ratio (BR) (Fig. 1D). Large BRs (closer to 1) represent less visualization of the target and more visualization of the neutral distractor (i.e., the greater the BR, the more difficult the task). The first trial of the titration phase was set to a BR of 0.25, increasing or decreasing by 0.05 following successful and unsuccessful trials, respectively. BR was restricted to the 0.15 to 0.90 range. At the end of the titration phase, target detection accuracy was calculated for each BR level, identifying the mode BR level within the 50-65% range. This level was then used for the entirety of the subsequent experimental phase. If a BR was not identified within this specified accuracy range, 0.05 above the mode BR was selected.

### Experiment 1 Procedure

The goal of experiment 1 was to validate our RSVP paradigm for studying EIB, ensuring results aligned with those of previously established approaches (Kennedy & Most, 2015; Most et al., 2005; Most & Jungé, 2008; Most & Wang, 2011). We hypothesized that negative distractors directly preceding the target, or probe distractors, would lead to impaired target detection, demonstrated by decreased accuracy in the choice phase. Experiment 1 consisted of 144 trials, with each image presented for a duration of 100 ms. We tested two probe distractor conditions, negative and scrambled negative, compared to a baseline neutral distractor. In the neutral condition, all images in the stream were neutral IAPS images. For the negative and scrambled negative conditions, one neutral distractor was replaced with either a scrambled negative or negative image. All images were pulled from the IAPS database, where negative images depicted emotionally salient conditions such as threatening animals, violence, or medical trauma (Bradley & Lang, 2017; Wang et al., 2012). Images in the negative condition were then used to create the scrambled negative images, following steps described in Most et al. (2005). The purpose of this scrambled negative condition was to ensure that observed negative effects are a result of valence rather than low-level visual features. In line with previous EIB designs, target stimuli appeared at the 2^nd^, 4^th^, or 8^th^ position within the sequence, with negative distractors preceding this target by 1, 2, or 8 positions. Probe distractor and lag conditions were randomized across trials.

### Experiment 2 Procedure

In experiment 2, we aimed to determine the extent to which image presentation duration modulates observed EIB phenomena. Experiment 2 consisted of 180 trials with images presented at a range of durations: 33.33, 50, 100, 150, 200, and 300 ms per image. Image duration conditions were tested using a block design, where each block consisted of 10 trials. Blocks were selected to be run in a pseudo-randomized fashion, without replacement, such that the same type of block was never repeated in immediate succession and all block types were run before any were repeated. Each type of block was run 3 times, totaling 30 trials for each image duration. Target stimuli appeared at the 3^rd^, 6^th^, 9^th^, or 12^th^ position within the sequence, with negative distractors preceding this target by 1 position (lag 1). Probe distractor conditions, neutral and negative, were randomized across trials.

Participants who completed experiment 2 also reported subjective image ratings following the experiment. Online software PsyToolkit was used for collecting ratings data (Stoet, 2010; Stoet, 2017). Participants rated all (22) negative images and a subset (28) of neutral images. Ratings were collected for both valence and arousal using a block design, with each rating condition presented in randomly ordered blocks. Within each block, images were also randomized. For the valence rating condition, participants were instructed to rate images on a scale from 1, unpleasant, to 7, pleasant. For the arousal rating condition, participants were instructed to rate images on a scale from 1, relaxed, to 7, stimulated.

### Experiment 3 Procedure

The objective of experiment 3 was to determine if the ability to recall a negative distractor was associated with EIB. Following the choice phase, participants performed a 5000 ms recall phase (Fig. 1E). During the recall phase, participants were shown 6-image options where they were instructed to select the image that appeared in the stream. Only one of the six options had been in the RSVP sequence, and this correct response was always the image that directly preceded the target (i.e., the lag 1 distractor). The options included an equal number of negative and neutral images, presented in a randomized order. Feedback was provided using the same procedure as in the choice phase.

Experiment 3 consisted of 160 trials with images presented at a range of durations—83.33, 100, 200, and 300 ms per image—following a randomized and non-consecutive design. Blocks were repeated four times, totaling 40 trials for each image duration. Target stimuli appeared at the 3^rd^, 6^th^, 9^th^, or 12^th^ position within the sequence, with negative distractors preceding this target by 1 position (lag 1). Probe distractor conditions, neutral and negative, were randomized across trials.

### Statistical Analyses

All analyses were performed using R Statistical Software in R Studio (RCoreTeam, 2024; RStudioTeam, 2020). Bayesian logistic regression models were fit using the R package ‘‘brms” to estimate the predictive power of task conditions in estimating accuracy in the choice phase (Bürkner, 2018, 2021; Kruschke, 2015). Specifically, Bayesian analysis was used to examine the proportion of posterior distributions above or below zero, assessing the magnitude by which task conditions influence the likelihood of accurate target detection during the choice phase (Kruschke, 2015).

All models were mixed effects models, allowing for estimation of group and individual parameters. A uniform prior was used for all models. Details regarding each model are included in the Appendix. For experiment 1, we modeled the effect of each type of probe distractor on accuracy at each lag, using neutral as the reference condition. For experiment 2, we estimated the effect of probe distractors on accuracy across image duration conditions, again using neutral as the reference condition. We also examined the influence of rating type (i.e., arousal and valence) on target accuracy. In experiment 3 we again estimated the effect of probe distractors on accuracy across each image duration, using neutral as the reference condition. We then created a model of target accuracy for each distractor condition (i.e., neutral and negative) with recall and image duration as weights. To assess substantial differences across model estimates, we compared posterior distributions by calculating the proportion of samples where one posterior distribution was meaningfully different from another, following established methods (McHugo et al., 2013; Olatunji, 2021; Onie & Most, 2020). In this context, we define ‘meaningful’ as instances where at least 95% of posterior samples are non-overlapping.

We conducted posterior predictive checks (PPCs) to assess how well the fitted model represents the observed data (Kruschke, 2015). This involved generating simulated data from the posterior distribution, using the “bayesplot” package in R, confirming that simulated data falls within the averaged observed response. We estimated the Bayes factor using the “brms” package in R to assess the strength of evidence for each model relative to a null model (Bürkner, 2018, 2021; Kruschke, 2015). Details regarding PPCs and our null model can be found in the Appendix.

## Results

### Experiment 1: Emotionally salient distractors immediately preceding the target impaired choice accuracy

We conducted experiment 1 to assess our modified RSVP design in the context of prior work. Participants were instructed to look for a bird image (the “target”) amongst a 17-image stream, in which probe distractors could appear at 1, 2, or 8 images preceding this target. Probe distractors were either neutral, negative, or scrambled negative. We hypothesized that negative probe distractors would be associated with decreased target accuracy in the choice phase. In line with previous work, we expected that effects of negative probe distractors on target accuracy would attenuate as a function of increasing lag (Kennedy & Most, 2011; Most et al., 2005).

We first compared target accuracies as a function of lag and probe distractor condition (Fig. 2A). For the neutral condition the mean target accuracy (± SEM) was 58.5% ± 1.52% (different “lags” in this case did not alter the image stream). For the negative condition, we estimated mean accuracies as 49.6% ± 2.68%, 45.5% ± 2.69%, and 59.8% ± 2.63% for lags 1, 2, and 8, respectively (Table A1). For the scrambled negative condition, we estimated mean accuracies as 57.4% ± 2.67%, 56.0% ± 2.66%, and 57.5% ± 2.65% for lags 1, 2, and 8, respectively. In summary, trials with negative probe distractors shown in closer proximity to the target demonstrated the largest accuracy decrements relative to neutral probe conditions.

**Figure 2.**
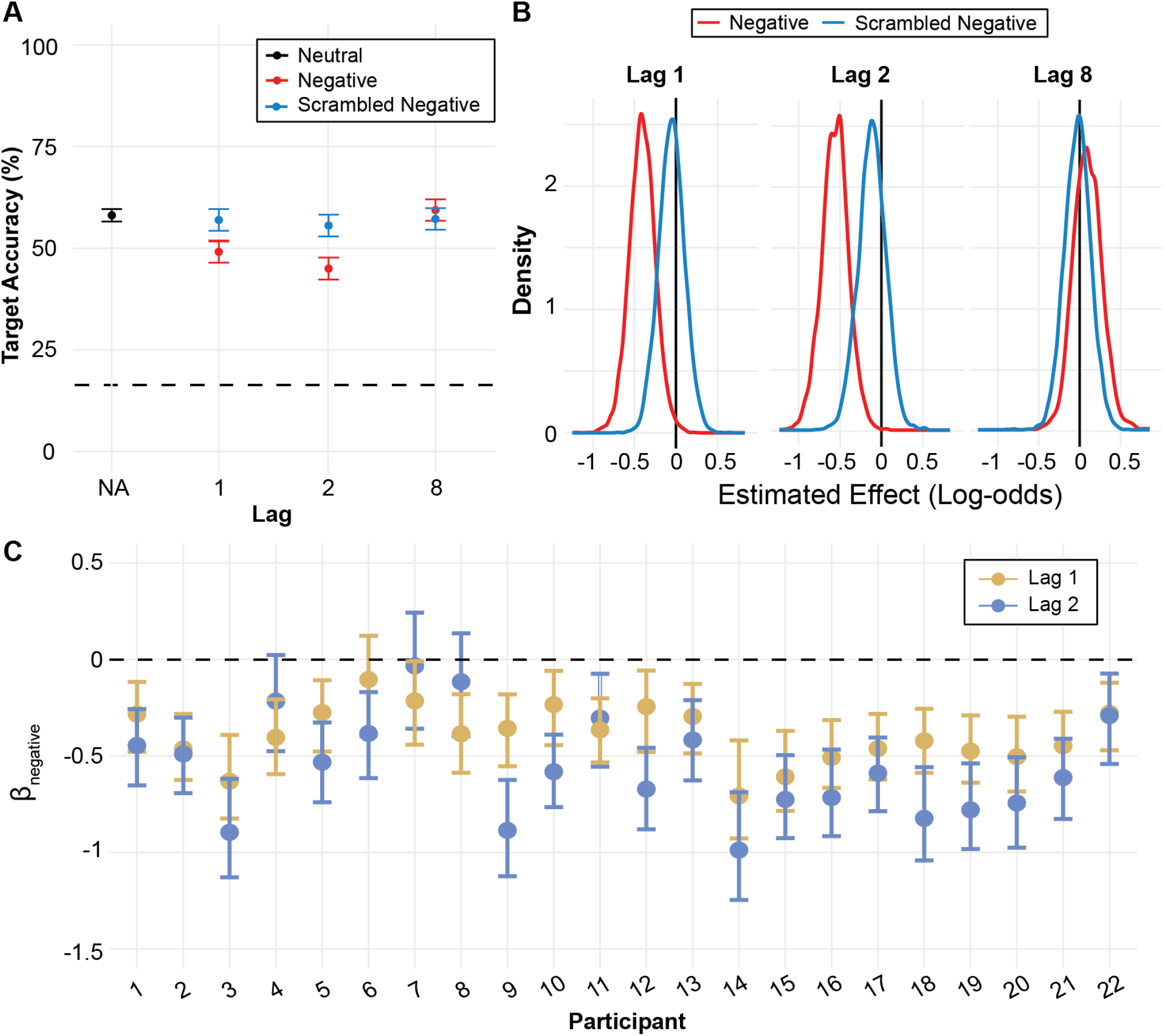
***A)***Target detection accuracy by distractor type and lag. Points and error bars represent mean and SEM, respectively. ***B)***Posterior distribution of estimated effect of distractor (scrambled negative—blue, negative—red) on accuracy. Negative and positive values indicate distractors worsen and improve accuracy, respectively. Left, middle, and right subplots show posterior distributions at lags 1, 2, and 8, respectively. ***C)***Participant-level weights for estimated effect of negative distractor on target accuracy at lags 1 (tan) and 2 (purple). Points and error bars represent mean and interquartile range (IQR), respectively. 17 of 22 participants show a larger effect of negative distractors at lag 2 than at lag 1.

We used Bayesian logistic regressions to model the effect of probe distractor condition on target accuracy at each lag position (Fig. 2B). We found no substantial differences in target accuracy between scrambled negative and neutral conditions. Comparing negative versus neutral distractor trials, we estimated effects of −0.39 (95% CI [−0.69, −0.09]), −0.56 (95% CI [−0.89, −0.24]), and 0.08 (95% CI [−0.24, 0.42]) for lags 1, 2, and 8, respectively, demonstrating an impairment in performance at shorter lags. Bayes factors (BF) were BF_10_<1, 30<BF_10_<100, and BF_10_<1 for estimated effects of negative distractor conditions on target accuracy at lags 1, 2, and 8, respectively. Only the BF estimate at lag 2 indicated overwhelming evidence in favor of modeled effects of probe distractor type on target accuracy.

To assess the robustness of group-level effects, we measured by-participant estimates of the negative distractor condition on accuracy at lag 1 and lag 2 (Fig. 2C). We observed numerically larger effects of negative probe distractors shown at lag 2 compared to lag 1 on both a group-and by-participant level. Specifically, 17 out of 22 participants showed a larger effect of trials with negative distractors at lag 2 than at lag 1. Posterior comparisons were then conducted to determine if estimated effects of distractor type across lags 1 and 2 were meaningfully different. We found that 76% of posterior samples from lags 1 and 2 were non-overlapping, indicating little meaningful difference in the effect of negative distractors on accuracy between lag 1 and lag 2. A summary of all posterior comparisons is included in Tables S2-3, with PPCs shown in Fig. A1.

### Experiment 2: Negative effects on target accuracy persisted with increasing image duration

Experiment 2 characterized the effect of image presentation duration on the EIB. Specifically, we tested EIB effects at the following image duration times: 33.33, 50, 100, 150, 200, and 300 ms. The presentation of a negative distractor directly preceding the target (lag 1) was randomized across trials compared to the baseline condition with all neutral distractors. We hypothesized that target choice performance would improve with increased image duration. Further, we predicted that the effect of negative probe distractors would decrease with increasing image duration.

First, we estimated target accuracy as a function of image duration, collapsing across probe distractor type. The mean accuracies (± SEM) were 17.4% ± 1.07%, 23.1% ± 1.19%, 46.4% ± 1.41%, 65.9% ± 1.34%, 71.7% ± 1.27%, and 78.1% ± 1.74% for 33.33, 50, 100, 150, 200, and 300 ms image durations, respectively (Fig. A2A), indicating target accuracy increases with longer image durations. Choice accuracy for the 33.33 ms and 50 ms image duration conditions were not significantly better than chance.

We used a Bayesian logistic regression model to estimate the group-level effects of image duration on accuracy, referenced to the 33.33 ms image duration (Fig. A2B). We estimated average effect sizes of 0.38 (95% CI [0.16, 0.59]), 1.47 (95% CI [1.25, 1.69]), 2.34 (95% CI [2.14, 2.56]), 2.64 (95% CI [2.43, 2.87]), and 2.96 (95% CI [2.64, 3.29]), for image durations of 50, 100, 150, 200, and 300 ms, respectively, indicating with 95% confidence that accuracy improved with increasing image duration. Effects of image duration on target accuracy were consistent on by-participant level (Fig. A2C). Comparisons of posterior distributions for each image duration revealed with 95% confidence that target accuracy improved with each duration increase. BF_10_>100 for image durations exceeding 50 ms, suggesting overwhelming evidence in favor of these modeled effects of image duration on target accuracy.

We next calculated target accuracy as a function of image duration and probe distractor type (Fig. 3A). For the negative distractor condition, mean accuracies (± SEM) were 18.0% ± 1.52%, 22.7% ± 1.67%, 39.6% ± 1.96%, 58.1% ± 1.97%, 64.4% ± 1.92%, and 71.5% ± 2.67% for 33.33, 50, 100, 150, 200, and 300 ms image durations, respectively (Table A4). For the neutral distractor condition, mean accuracies (± SEM) were 16.8% ± 1.49%, 23.6% ± 1.69%, 53.3% ± 1.99%, 73.8% ± 1.76%, 79.1% ± 1.63%, and 84.6% ± 2.14% for 33.33, 50, 100, 150, 200, and 300 ms image durations, respectively. In summary, trials with negative probe distractors demonstrated consistently numerically lower target accuracies across image durations.

**Figure 3.**
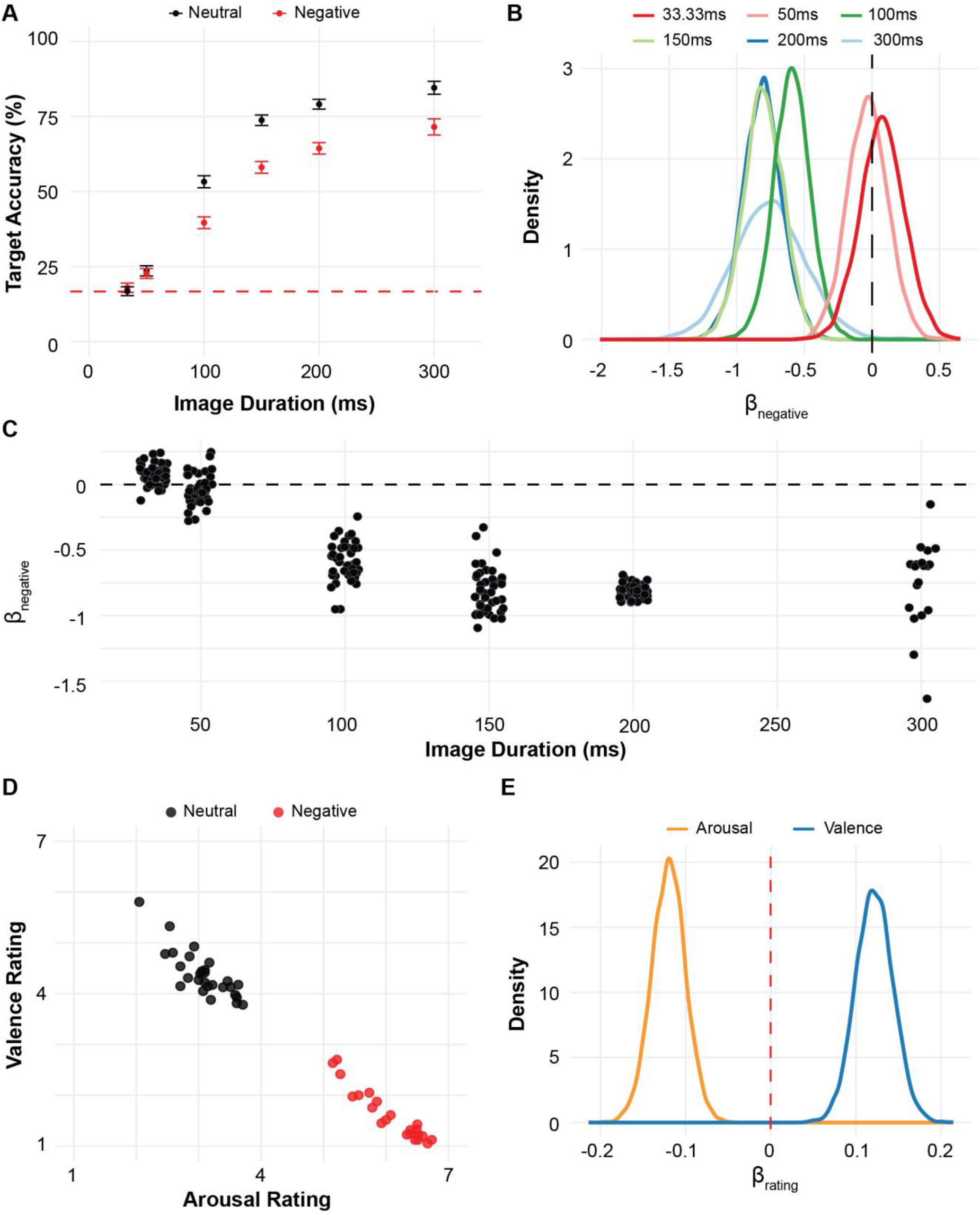
***A)***Target detection accuracy as a function of image duration by distractor type. Point and error bar represent mean ± SEM. ***B)*** Group-level weights for estimated effect of negative distractor on target accuracy at each image duration, referenced to the neutral distractor condition. Black dashed line represents zero-line where below zero represents worse performance for negative distractors. ***C)***Participant-level weights for estimated effect of negative relative to neutral probe distractor on accuracy at each image duration. Black dashed line represents zero-line where below zero represents worse performance for negative distractors. ***D)***Subjective rating of valence vs. arousal. Dots represent average rating for one IAPS image, where dots are colored by IAPS categorization. ***E)***Weights for estimated effect of rating type (i.e. arousal vs. valence) on accuracy. Red dashed line represents zero-line, where values above zero represent higher ratings corresponding to improved accuracy.

We used a Bayesian logistic regression model to estimate the effect of negative probe distractors on target accuracy at each image duration (Fig. 3B). Average effect sizes were 0.08 (95% CI [−0.25, 0.40]), −0.03 (95% CI [−0.33, 0.26]), −0.59 (95% CI [−0.87, −0.33]), −0.80 (95% CI [−1.10, −0.53]), −0.81 (95% CI [−1.10, −0.52]), and − 0.76 (95% CI [−1.27, −0.23]) at image durations of 33.33, 50, 100, 150, 200, and 300 ms, respectively, indicating with 95% confidence that the trials with negative relative to neutral probe distractors impaired target accuracy for image durations exceeding 50 ms (Table A5). Observed effects of negative distractor conditions on target accuracy held up on a by-participant level, with slightly higher variance of estimated effects at image duration 300 ms (Fig. 3C). Comparisons of posteriors for these conditions revealed no meaningful differences for negative condition effects across image durations ranging from 100 to 300 ms (Table A6). BF_10_>100 for image durations exceeding 50 ms, indicating overwhelming evidence in favor of modeled effects of distractor type on target accuracy at each image duration.

Finally, we assessed the relationship between subjective ratings of valence and arousal on target detection accuracy. Participants were instructed to rate the valence and arousal level of all negative images and an equal subset of neutral images (selected from neutral categorized images in the IAPS database). We observed that IAPS images categorized as neutral with low arousal were rated as such (Fig. 3D). This was the same for IAPS images categorized as negative with high arousal. For the trials with images that were rated, we modeled the relationship between rating and accuracy, determining estimated effects of 0.12 (95% CI [0.08, 0.16]) and −0.12 (95% CI [−0.16, −0.08]) for valence and arousal, respectively (Fig. 3E), consistent with the previously established notion that EIB varies in proportion to these subjective image ratings (Onie & Most, 2020). BF_10_>100 for both valence and arousal, demonstrating overwhelming evidence in favor of modeled effects of subjective ratings on target accuracy.

### Experiment 3: Probe distractor recall is associated with target accuracy for negative but not neutral trials

In experiment 3 we estimated the degree to which conscious perception or recollection of probe distractor images was associated with target detection. Experiment 3 closely mirrored experiment 2, with two key differences. The first being the set of included image duration conditions: 83.33, 100, 200, and 300 ms. More notably, experiment 3 also included a recall phase where participants were asked which image they recall appearing in the stream. We hypothesized that recall accuracy would be higher for negative compared to neutral probe distractors. Further, we hypothesized that recall would serve as a predictor of worse target accuracy on negative trials.

We first calculated target accuracy by image duration, collapsing across probe distractor type (Fig. A4A). The mean accuracies (± SEM) were 39.1% ± 1.44%, 49.0% ± 1.48%, 73.6% ± 1.31%, and 81.0% ± 1.17% for 83.33, 100, 200, and 300 ms image durations, respectively. We used a Bayesian logistic regression model to estimate the group-level effects of image duration on accuracy, referenced to the 83.33 ms image duration (Fig. A4B). We estimated average effect sizes of 0.42 (95% CI [0.24, 0.60]), 1.54 (95% CI [1.34, 1.75]), and 1.95 (95% CI [1.75, 2.16]), for image durations of 100, 200, and 300 ms respectively (BF_10_>100). Similar to experiment 2, descriptive statistics and Bayesian estimates demonstrated that accuracy improved with increased image duration. Effects of image duration on target accuracy were again consistent on a by-participant level (Fig. A4C). Comparisons of posterior distributions revealed that all posteriors were non-overlapping with 95% confidence, suggesting again that accuracy meaningfully improved with each duration increase.

We next calculated target accuracy by image duration and probe distractor type (Fig. 4A). For the negative distractor condition, mean accuracies (± SEM) were 35.0% ± 2.00%, 42.0% ± 2.06%, 69.0% ± 1.94%, and 78.8% ± 1.72% for 83.33, 100, 200, and 300 ms image durations, respectively (Table A7). For the neutral distractor condition, mean accuracies (± SEM) were 43.3% ± 2.08%, 56.1% ± 2.08%, 78.1% ± 1.73%, and 83.2% ± 1.57% for 83.33, 100, 200, and 300 ms image durations, respectively. In summary, negative distractor conditions demonstrated consistently numerically lower target accuracy than neutral distractor conditions across image durations, consistent with Experiment 2.

We used a Bayesian logistic regression model to estimate the predictive power of trials with negative probe distractors on accuracy at each image duration (Fig. 4B). Estimated model weights for the negative distractor condition were −0.38 (95% CI [−0.65, −0.11]), −0.60 (95% CI [−0.85, −0.35]), −0.50 (95% CI [−0.82, −0.20]), and − 0.28 (95% CI [−0.63, 0.06]) at image durations of 83.33, 100, 200, and 300 ms, respectively, indicating substantial impairments of trials with negative distractors on accuracy at all image duration conditions below 300 ms (Table A8). Observed effects of negative distractor conditions on target accuracy held up on a by-participant level (Fig. 3C). BFs were 1<BF_10_<3, BF_10_>100, 10<BF_10_<30, and BF_10_<1 for estimates of the negative distractor condition on target accuracy at image durations of 83.33, 100, 200, and 300, respectively. BF estimates indicate overwhelming evidence in favor modeled effects of trials with negative distractors on target accuracy at durations 100 and 200 ms, with weak evidence for this model at 83.33 ms.

Most notably, we examined the relationship between distractor recall and the probability of accurate target detection. We observed numerically lower target accuracies on trials where the negative probe distractor was successfully recalled (Fig. 4D). Table 2 includes a summary of target accuracy differences across image duration, recall, and distractor conditions. We then used logistic regression to model the degree to which image duration and distractor recall contribute to target accuracy for negative and neutral distractor conditions, respectively. For the neutral condition, image duration, recall, and interaction weights were estimated as 0.71 (95% CI [0.57, 0.86]), 0.05 (95% CI [−0.17, 0.27]), and 0.26 (95% CI [0, 0.51]), respectively. For the negative condition, these weights were estimated as 0.86 (95% CI [0.70, 1.02]), −0.42 (95% CI [−0.64, −0.20]), and 0.05 (95% CI [−0.17, 0.27]). BF_10_>100 for both neutral and negative models of target accuracy, demonstrating overwhelming evidence that recall of negative distractors, specifically, was associated with decreased target accuracy. On trials where no recall occurred, the selection of the correct distractor category (negative or neutral) was no better than chance, with participants more frequently selecting neutral valence distractors (Fig. A5). In summary, for the neutral distractor condition we estimated with 95% confidence an effect of rate, but not distractor recall, on target accuracy (Fig. 4E). Separately, for the negative distractor condition we estimated with 95% confidence an effect of both rate and recall on target accuracy.

**Figure 4.**
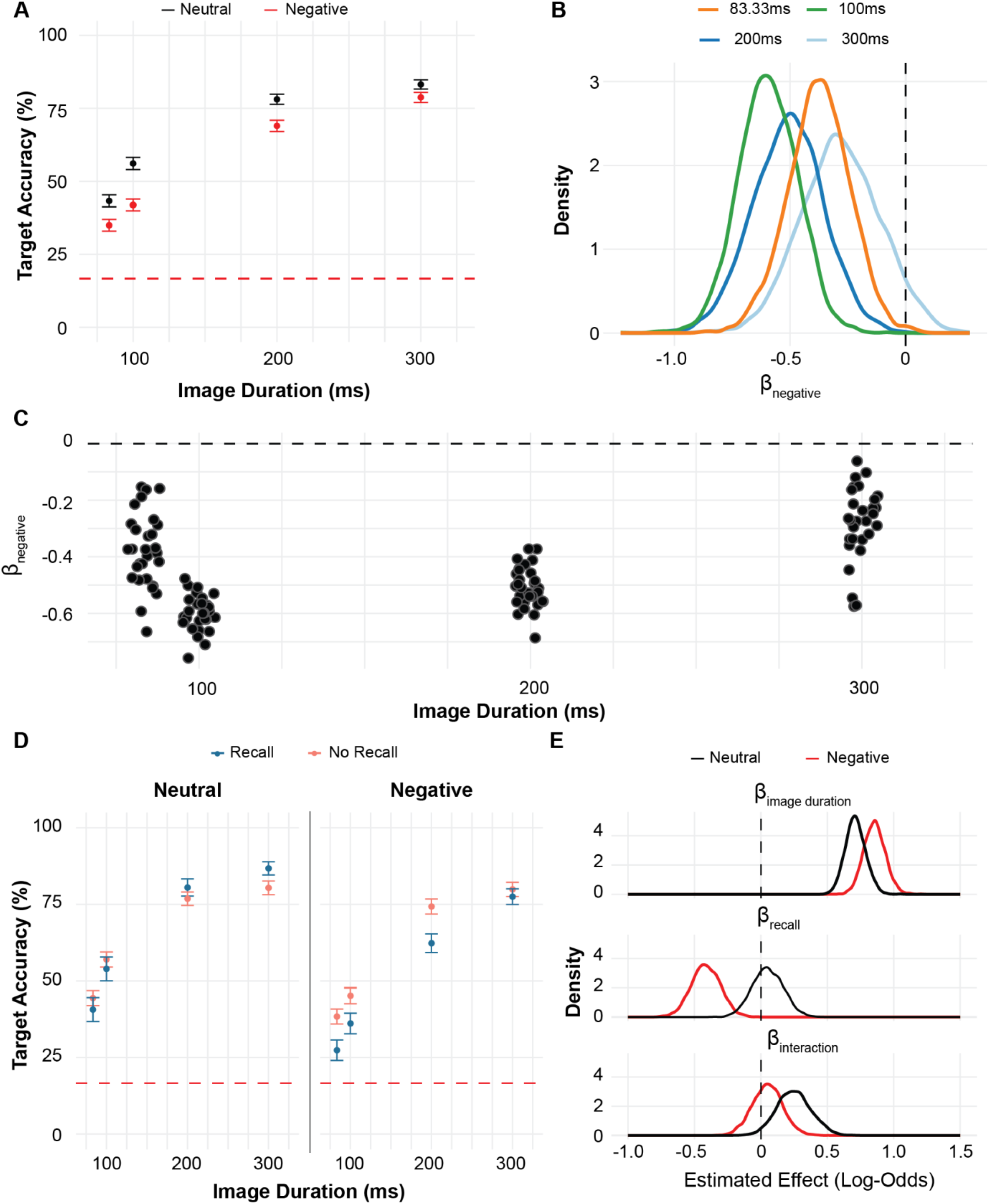
***A)***Target accuracy as a function of image duration by distractor type. Point and error bar represent mean ± SEM. ***B)***Weights for estimated group-level effect of negative distractor on target accuracy at each image duration, referenced to the neutral distractor condition. Black dashed line represents zero-line where below zero represents worse performance for negative distractors. ***C)***Participant-level weights for estimated effect of negative relative to neutral probe distractor on accuracy at each image duration. Black dashed line represents zero-line where below zero represents worse performance for negative distractors. ***D)***Target accuracy as a function of image duration by distractor recall. Subplots are for each distractor type. Point and error bar represent mean ± SEM. ***E)***Model weights for each predictor—image duration, distractor recall, and interaction between image duration and recall—using a Bayesian hierarchical logistic regression by distractor condition.

**Table 2.** Average target accuracy (%) by distractor, recall, and image duration conditions.

| Image Duration (ms) | Average Target Accuracy (%) |  |  |  |
| --- | --- | --- | --- | --- |
|  | Negative |  | Neutral |  |
|  | Recall | No Recall | Recall | No Recall |
| 83.33 | 27.4 | 38.4 | 40.6 | 44.4 |
| 100 | 36.1 | 45.1 | 54.0 | 57.0 |
| 200 | 62.3 | 74.3 | 80.5 | 76.8 |
| 300 | 77.5 | 79.9 | 86.7 | 80.3 |

## Discussion

Our study aimed to further characterize the conditions in which affectively charged stimuli impair visual attention. We conducted experiment 1 to validate our modified RSVP paradigm, confirming the presence of an EIB effect. In experiment 2, we expanded on the established temporal dynamics of EIB, focusing on the contribution of image presentation duration to the effect of negative distractors on target accuracy and assessing parametric influences of emotional distractors on performance. Finally, we conducted experiment 3 to assess the effect of distractor recall on accuracy in choice phase, to investigate whether there was an association between depth of processing and EIB. Together, these findings contribute to our understanding of how emotional stimuli impair cognitive processes, such as attention.

We conducted experiment 1 to establish that our updated RSVP design captured previously reported emotion induced blindness phenomena. In line with this work, we hypothesized that the presence of negative relative to neutral distractors would impair target detection, demonstrated by decreased accuracy during the choice phase. We also tested a scrambled negative condition, to confirm that negative impairments in target detection were due to valence rather than low-level visual features. Negative and scrambled negative distractor conditions were tested at three positions preceding the target: lags 1, 2, and 8. In line with previously established results, we demonstrated that negative relative to neutral distractor conditions led to significant impairments in accuracy.

Further, we confirmed that negative impairments in target accuracy attenuated with increased distance (lag) from the target, with no significant effects of negative distractors when shown 8 images preceding the target. Finally, we found no significant differences between scrambled negative and neutral distractor conditions, consistent with the notion that negative-distractor effects were not due to low-level visual features.

The goal of experiment 2 was to expand upon previously identified temporal dynamics of EIB. In particular, we examined the effect of image duration in modulating the EIB phenomena. We first demonstrated that target detection, collapsing across distractor conditions, improved with increased image duration. Further, we found that performance was not significantly better than chance at image durations of 33.33 and 50 ms. Next, we examined target accuracy as a function of both image duration and distractor condition. We determined with 95% confidence that performance on negative distractor trials was worse than neutral distractor trials across all image durations where performance was greater than chance. We conducted further analyses to determine if EIB effects observed at these durations were meaningfully different. We did not find any meaningful differences, suggesting that negative distractor trials demonstrate persistent effects on target detection when image durations exceed 50 ms.

Findings from experiment 2 regarding faster image presentations were unsurprising, as image duration directly modulates task difficulty. As such, the absence of EIB at faster presentation rates likely reflects a general inability to attend to individual stream images, including negative probe distractors. However, it was somewhat surprising that increased image duration did not modulate EIB effects. Given prior studies on temporal dynamics showing that increased lag attenuates EIB effects, one might have anticipated a similar pattern for longer image presentation durations. This attenuation did not occur, possibly because while increasing image duration permits more time to detect the target, it also enhances processing of emotional stimuli.

Experiment 3 further highlighted this tradeoff in which longer image durations aided target detection while, for some individuals, sustaining EIB effects. Although some participants showed persistent EIB at 300 ms, there was no significant group-level effect of trials with negative distractors. Therefore, while there is some low level persistent effect of EIB, it is possible that the degree of EIB with increasing image duration reflects individual differences with respect to anxiety and rumination (Kennedy & Most, 2015). In particular, those with higher anxiety and rumination characteristics have been reported to exhibit higher sensitivity to negative stimuli, shown by larger EIB at earlier lags. Therefore, one may hypothesize that inconsistent EIB effects at the 300 ms reflect individual variation in those features, where those with stronger EIB effects may tend towards higher levels of anxiety and rumination when shown negative images for extended periods.

We also implemented image ratings in experiment 2 to observe how subjective reports of valence and arousal might predict target accuracy. We determined with 95% confidence that lower valence (i.e., negative) and higher arousal (i.e., more stimulating) ratings were predictive of decreased target accuracy. Specifically, images perceived as more intensely negative led to greater impairments in subsequent target detection, consistent with previous findings that EIB is sensitive to gradations in valence and arousal (Onie & Most, 2020). This is in line with one previous study that has identified that EIB is sensitive to gradations in valence and arousal (Onie & Most, 2020). Identical to this work, this finding suggests that EIB effects arise from perceived valence and intensity of probe distractors rather than low-level image features.

Our aim in introducing the recall phase in experiment 3 was to assess the extent to which conscious perception of distractors and/or depth of processing contributes to target accuracy. We observed that target accuracy was worse for the negative relative to neutral distractor condition (Fig. 4D), regardless of recall. However, successful recall of negative distractors also served as a predictor of worsened target accuracy, which was not the case for neutral distractors. Observed relationships between negative distractors, recall, and target accuracy further suggest that EIB is a graded phenomenon, where emotionally salient stimuli processed deeply enough for recall will produce the largest impairment on attention. This expands on recent work indicating that EIB itself is not strictly an all-or-none impairment (Keefe & Zald, 2022). Rather than recall of probe distractors, it was determined that target processing following distractors occurs on a continuum, with participants reporting varying levels of awareness of the target image following exposure to negative and neutral distractors (Keefe & Zald, 2022). Our findings further support this idea, suggesting conscious perception of negative distractors also operates on a graded scale rather than an all-or-none mechanism in producing EIB effects.

We also examined trials where subjects failed to correctly recall the distractor. As recall choices included equal numbers of negative and neutral stimuli, we aimed to identify if trials with inaccurate recall of the exact distractor item nonetheless demonstrated accurate recall of distractor valence. Specifically, in negative distractor trials, we sought to understand whether participants were generally aware that something negative appeared in the stream, even if they could not precisely identify it. However, such a pattern was not evident, and there was a general bias to select neutral images when participants were unable to recall the exact distractor item (Fig. A5). This bias was perhaps due to vigilance-avoidance mechanisms, where avoiding potentially threatening stimuli may be adaptive (Goodhew & Edwards, 2022; Proud et al., 2020).

Several theoretical models have been proposed to explain EIB, most notably the two-stage and spatiotemporal competition accounts (Chun & Potter, 1995; Desimone, 1998; Desimone & Duncan, 1995; Wang et al., 2012). The two-stage model suggests that stimuli are initially detected (stage 1) before undergoing consolidation (Chun & Potter, 1995; Potter et al., 2002). However, this consolidation process is capacity-limited, creating a bottleneck that restricts how many stimuli can be encoded at a given time. Therefore, stimuli in stage 1 that are waiting for available processing capacity in stage 2 risk being overwritten by subsequent stimuli. If EIB were solely due to the limited capacity of late-stage consolidation, we would expect longer image durations to significantly mitigate EIB. However, while prolonged image duration improved overall performance, some persistent EIB effects remained. This persistence contradicts a purely late-stage explanation, suggesting that earlier-stage mechanisms may also play a role in EIB.

The spatiotemporal competition account provides an alternative explanation for observed EIB phenomena (Desimone & Duncan, 1995; Most & Wang, 2011; Wang et al., 2012). Based on models of biased competition, this account suggests that visual stimuli that appear close in space and time compete for neural representation of a shared receptive field (Desimone & Duncan, 1995; Kennedy et al., 2018; Wang et al., 2012). Unlike the two-stage model, which focuses on attentional bottlenecks during later-stage consolidation of stimuli, the spatiotemporal competition model suggests that targets and emotional distractors may compete for earlier-stage detection of stimuli. Most notably, biased competition accounts for the observed graded nature of EIB, where attentional allocation is distributed between targets and probe distractors based on their relative salience. In particular, our findings demonstrate that the extent to which attention is diverted from targets depends on several factors: exposure to negative distractors, the degree to which these distractors are processed, and their perceived emotional intensity. Each of these factors increases the likelihood that probe distractors will be allocated more attentional resources, thereby impairing target detection.

In this study, we aimed to characterize the conditions necessary to elicit EIB. Our findings from experiment 1 reinforced the robust impact of emotionally salient stimuli on attention, where negative-valence distractors impaired subsequent target detection. Results from experiments 2 and 3 indicate the persistent effects of EIB despite increasing image duration, suggesting that effects of EIB are not purely due to capacity limitations on later-stage image encoding. Experiment 2 revealed that EIB is modulated by subjective ratings of valence and arousal, with more intensely negative images producing larger attentional impairments. From experiment 3, we observed that the depth of processing of negative distractors also modulates EIB, where probe distractors recalled produced larger impairments on subsequent target performance. Collectively, these findings support the notion that EIB is a graded rather than an all-or-nothing phenomenon, modulated by exposure, perceived negative intensity, and depth of processing.

## Acknowledgements

We are grateful for the guidance of Dr. Guillaume Pagnier on statistical analysis procedures

## Data Availability

De-identified datasets for the current study are available from the corresponding author on reasonable request.

## Appendix

### Methods: Models by Experiment

We used Bayesian hierarchical logistic regression models to estimate the effect of predictors from each experiment on target accuracy. Models were fit using the R package ‘‘brms,” where each included the fixed and random effects. Details on each model by experiment are included below. Bayes factor was estimated by comparing each model to a null model, using the following:

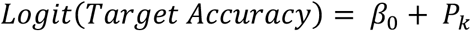

*P*_*k*_ represents the random effects for each participant, allowing model estimates on a by-participant level. The estimated effects (fixed and random) are expressed as log-odds weights, which describe the effect of predictors on the probability of a correct target response. *β*_0_ represents the baseline log-odds target accuracy, assuming no effects of other predictors on target accuracy.

*Experiment 1.* In experiment 1 we examined the effect of distractor type on target accuracy at each lag. As such, we ran three models for lags 1, 2, and 8, using the following:

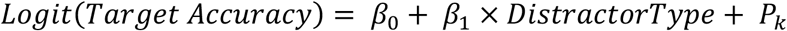

In this case, *β*_0_ represents the baseline log-odds target accuracy when all stream images are neutral (i.e., neutral distractor condition). *β*_1_ is the weight associated with distractor type, where values > 0 indicate the presence of that distractor type is associated with a higher probability of selecting the correct target.

*Experiment 2.* In experiment 2 we first examined the effect of image duration on target accuracy, using the following logistic regression model:

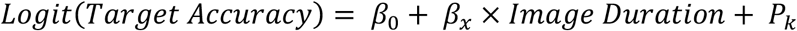

In this case, *β*_0_ represents the log-odds accuracy for the reference level, 33.33 ms. *β*_*x*_ captures the log-odds change in accuracy for each rate level (i.e., *β*_100_, *β*_150_, etc.). Image duration is treated as an ordered factor with levels for each rate category (i.e., 33.33, 50, 100, etc.).

Next, we examined the effect of distractor type on target accuracy at each image duration. As such, we ran models for each image duration category (6 total), using the following:

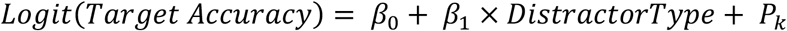

In this case, *β*_0_ represents the baseline log-odds target accuracy when all stream images are neutral (i.e., neutral distractor condition). *β*_1_ is the weight associated with distractor type, where values > 0 indicate the presence of that distractor type is associated with a higher probability of selecting the correct target.

*Experiment 3.* We repeated models from experiment 2 to compare target accuracies across rates both by and collapsing distractors. Next, we estimated the relative contribution of image duration and recall by distractor condition. This resulted in two models, shown below:

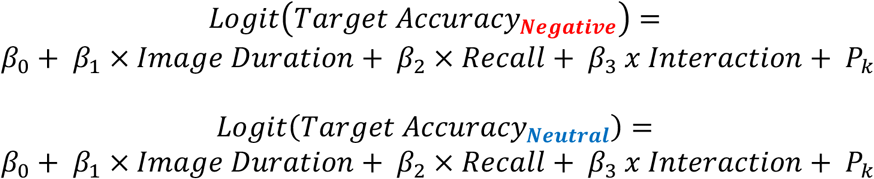

*β*_0_ represents the baseline log-odds target accuracy when all stream images are neutral (i.e., neutral distractor condition). *β*_1_ is the weight associated with image duration, where values > 0 indicate the longer image durations are associated with a higher probability of selecting the correct target. *β*_2_ is the weight associated with recall of distractor images, where values > 0 indicate accurate distractor recall is associated with a higher probability of selecting the correct target. Finally, *β*_3_ represents the interaction between recall and image duration, where values capture how the effect of image duration on target accuracy might change depending on recall accuracy.

## Supplementary Results

**Table A1.** Average target accuracy (%) by distractor and lag conditions.

| | Average (%) Target Accuracy (± SEM) | | |
| --- | --- | --- | --- |
| Lag | Neutral | Negative | Scrambled Negative |
| NA | 58.5 ± 1.53 | | |
| 1 | | 49.6 ± 2.68 | 57.4 ± 2.67 |
| 2 | | 45.5 ± 2.69 | 56.0 ± 2.66 |
| 8 | | 59.8 ± 2.63 | 57.6 ± 2.65 |

**Table A2.** Model Estimates of Distractor Type Effects on Target Accuracy.

|  | Model Weights (Lower CI, Upper CI) |  |
| --- | --- | --- |
| Lag | Negative | Scrambled Negative |
| 1 | -0.39 (-0.69, -0.09) | -0.05 (-0.34, 0.23) |
| 2 | -0.56 (-0.89 -0.24) | -0.10 (-0.41, 0.23) |
| 8 | 0.08 (-0.24, 0.42) | -0.03 (-0.33, 0.28) |

**Table A3.** Experiment 1 Model Comparisons of Distractor Type Effects on Target Accuracy.

|  | Proportion of Non-Overlapping Posterior Values |  |
| --- | --- | --- |
| Comparison | Negative | Scrambled Negative |
| Lag 1 > Lag 2 | 0.756 | 0.396 |
| Lag 2 > Lag 8 | 0.003 | 0.617 |
| Lag 1 > Lag 8 | 0.018 | 0.533 |

**Figure A1.**
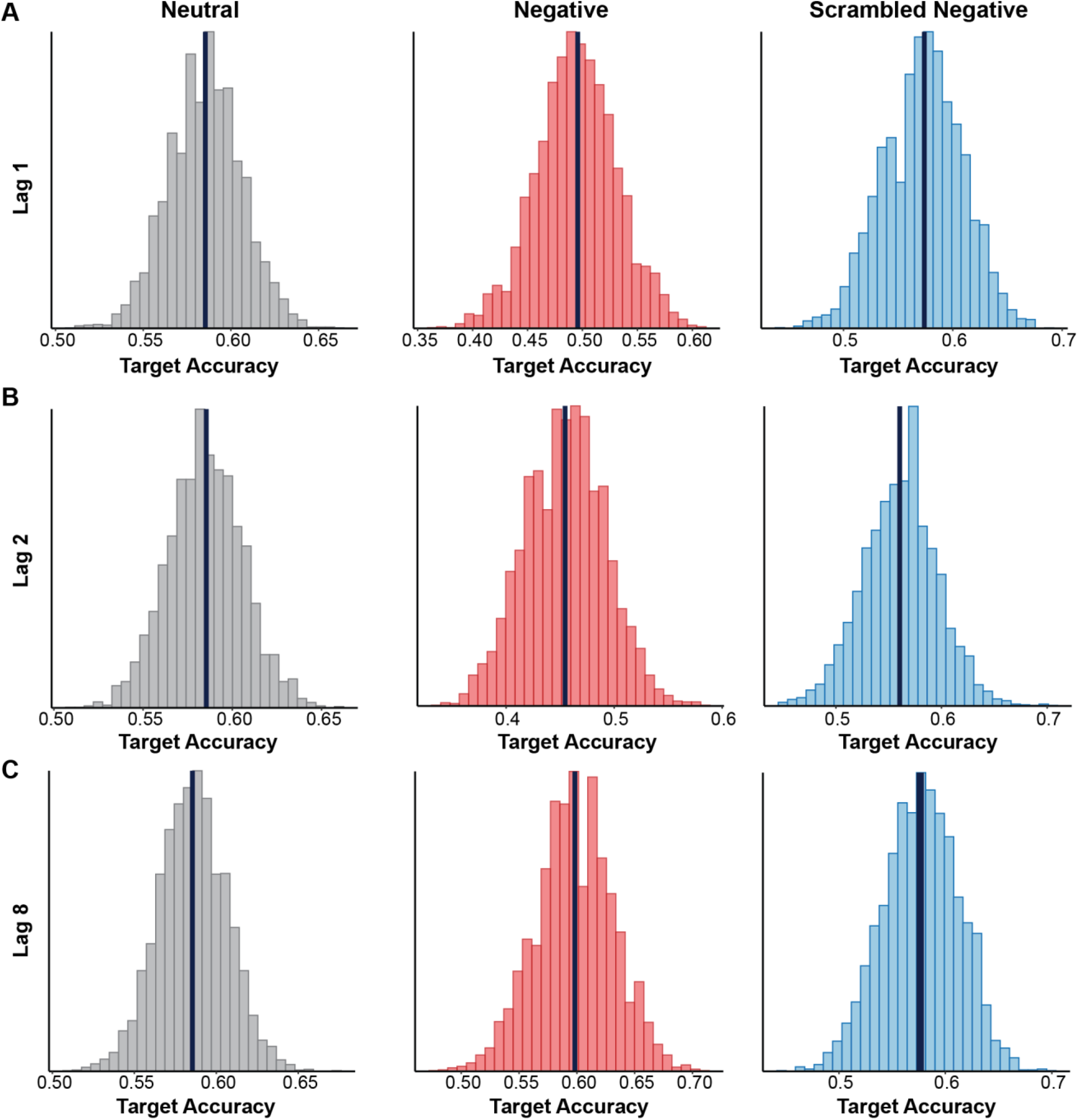
Experiment 1 observed average (black vertical lines) and simulated distribution (histogram) target accuracies for neutral (gray), negative (red), and scrambled negative (blue) conditions. ***A)***Effect of distractor type on target accuracy at lag 1. ***B)***Effect of distractor type on target accuracy at lag 2. ***C)***Effect of distractor type on target accuracy at lag 8.

**Figure A2.**
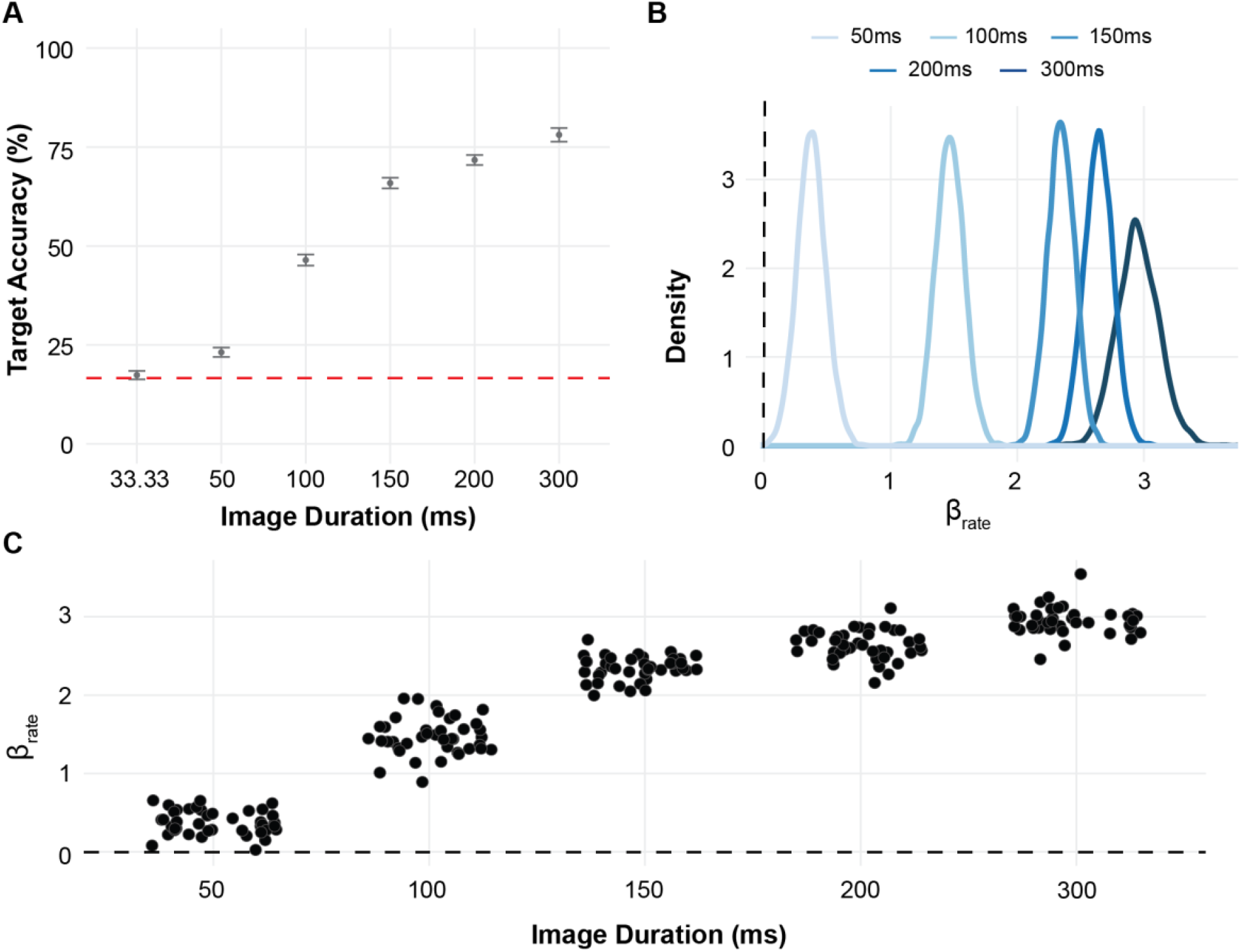
Experiment 2 ***A)***target accuracy as a function of image presentation duration (ms). Point and error bar represents mean ± SEM. Dashed red line represents chance performance at 16.67%. ***B)***Group-level estimated effect of image duration on accuracy, referenced to the 33.33ms image duration. Black dashed line represents zero-line, where values above zero indicate increased rate improves accuracy. ***C)***Participant-level estimated effect of image duration on accuracy, referenced to the 33.33ms image duration. Black dashed line represents zero-line, where values above zero indicate increased rate improves accuracy.

**Table A4.** Average target accuracy (%) by distractor and image duration.

| | Average (%) Target Accuracy (± SEM) | | | | | |
| --- | --- | --- | --- | --- | --- | --- |
|  | 33.33 | 50 | 100 | 150 | 200 | 300 |
| Neutral | 16.8 ± 1.49 | 23.6 ± 1.69 | 53.3 ± 1.99 | 73.8 ± 1.76 | 79.1 ± 1.63 | 84.6 ± 2.14 |
| Negative | 18.0 ± 1.52 | 22.7 ± 1.67 | 39.6 ± 1.96 | 58.1 ± 1.97 | 64.3 ± 1.92 | 71.5 ± 2.70 |

**Table A5.** Estimated effect of negative distractors by image duration.

| Image Duration (ms) | Model Weight (Lower CI, Upper CI) |
| --- | --- |
| 33.33 | 0.08 (-0.25, 0.40) |
| 50 | -0.03 (-0.33, 0.26) |
| 100 | -0.59 (-0.87, -0.33) |
| 150 | -0.80 (-1.10, -0.53) |
| 200 | -0.81 (-1.10, -0.52) |
| 300 | -0.76 (-1.27, -0.23) |

**Table A6.** Model comparisons for estimated effects of negative distractors by image duration.

| Comparison | Proportion of Non-Overlapping Posterior Values |
| --- | --- |
| 100 > 150 | 0.853 |
| 150 > 200 | 0.513 |
| 200 > 300 | 0.422 |
| 100 > 200 | 0.868 |
| 100 > 300 | 0.717 |
| 150 > 300 | 0.428 |

**Figure A3.**
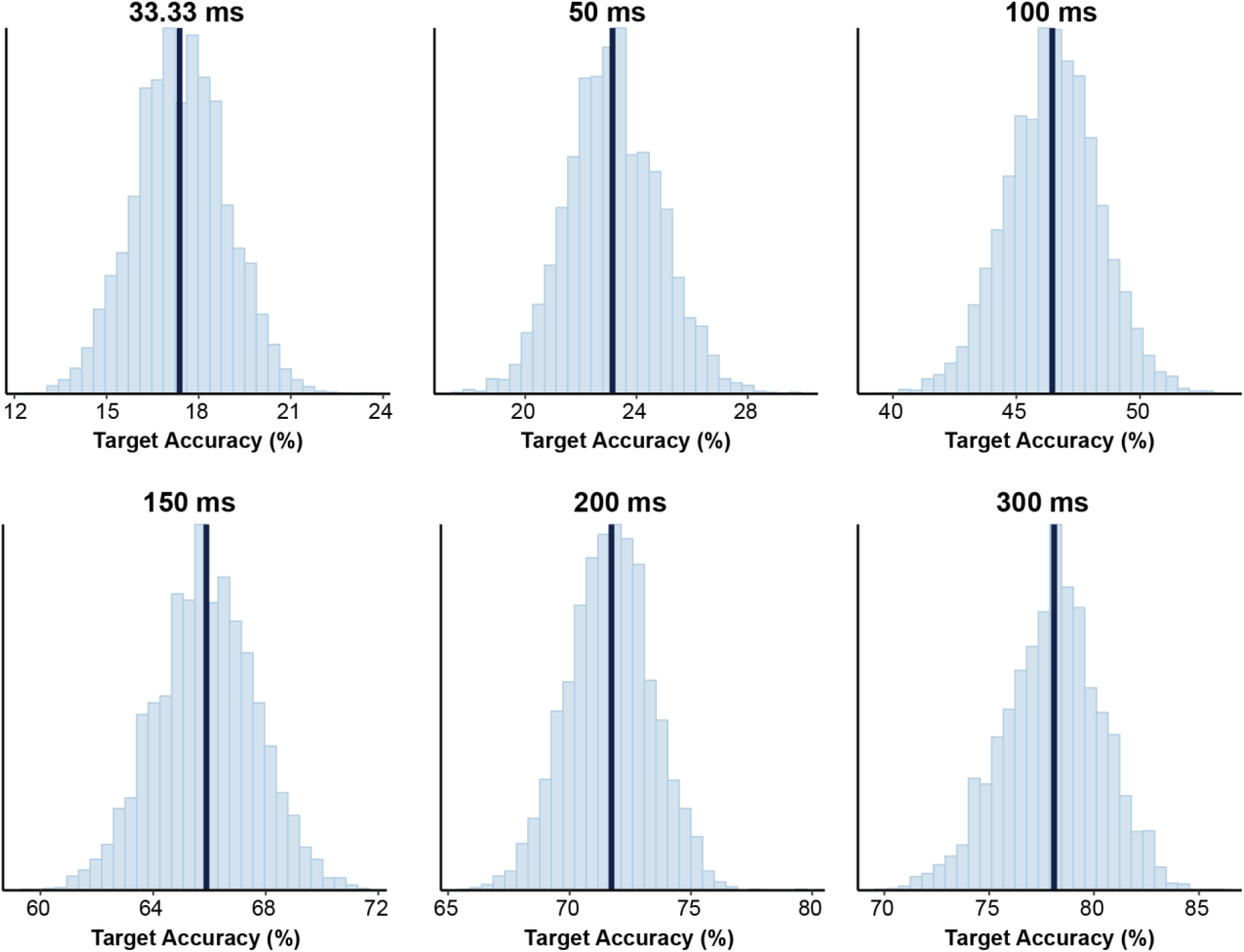
Experiment 2 observed average (black vertical lines) and simulated distribution (histogram) target accuracies for each image duration. Simulated data were created using model weights for estimating effect of negative distractors on target accuracy at each image duration condition.

**Figure A4.**
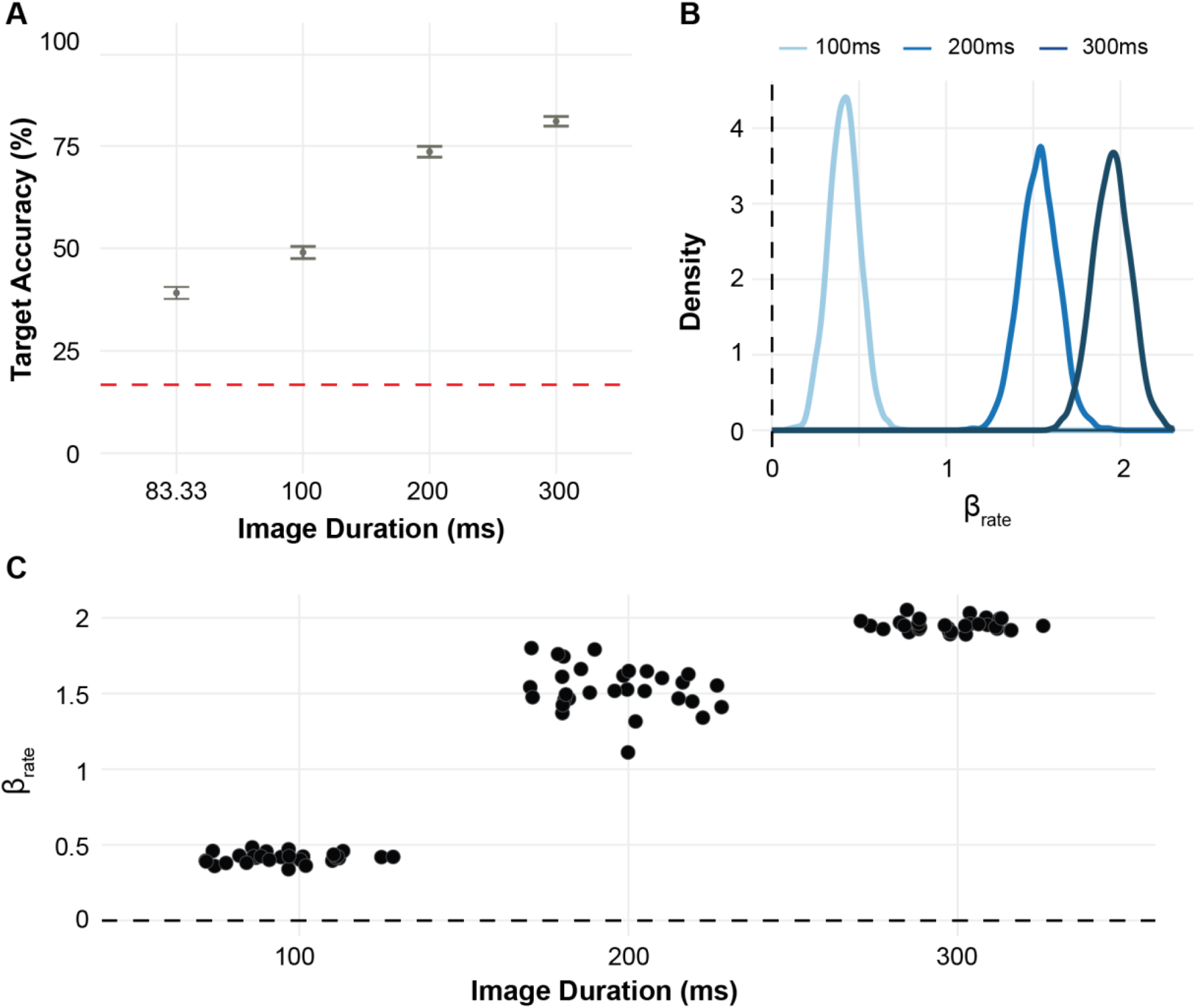
Experiment 3 ***A)*** target accuracy as a function of image presentation duration (ms). Point and error bar represents mean ± SEM. Dashed red line represents chance performance at 16.67%. ***B)*** Group-level estimated effect of image duration on accuracy, referenced to the 83.33ms image duration. Black dashed line represents zero-line, where values above zero indicate increased rate improves accuracy. ***C)*** Participant-level estimated effect of image duration on accuracy, referenced to the 83.33ms image duration. Black dashed line represents zero-line, where values above zero indicate increased rate improves accuracy.

**Table A7.** Average target accuracy (%) by distractor and image duration.

| | Average (%) Target Accuracy (± SEM) | | | |
| --- | --- | --- | --- | --- |
|  | 83.33 | 100 | 200 | 300 |
| Neutral | 43.3 ± 2.08 | 56.1 ± 2.08 | 78.1 ± 1.73 | 83.2 ± 1.57 |
| Negative | 35.0 ± 2.00 | 41.9 ± 2.06 | 69.0 ± 1.94 | 78.8 ± 1.72 |

**Table A8.**
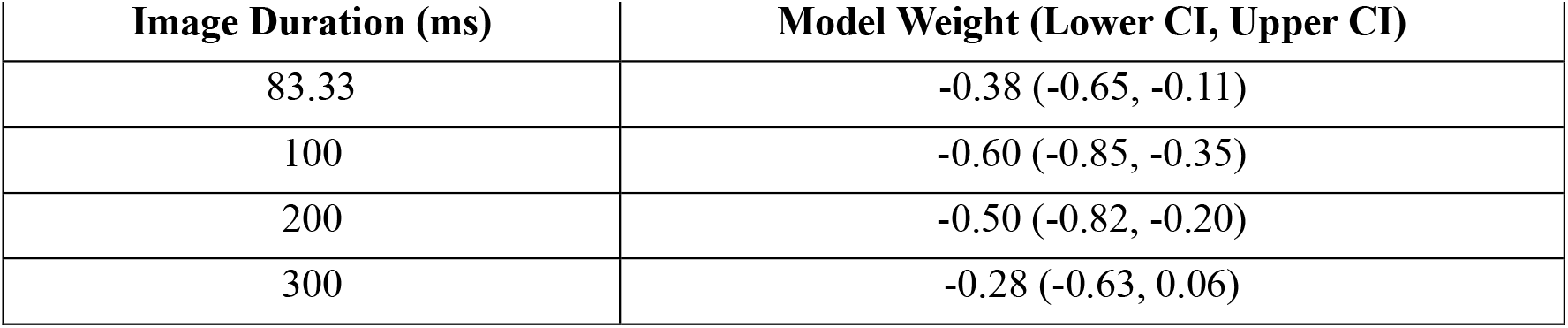
Estimated effect of negative distractors by image duration.

**Figure A5.**
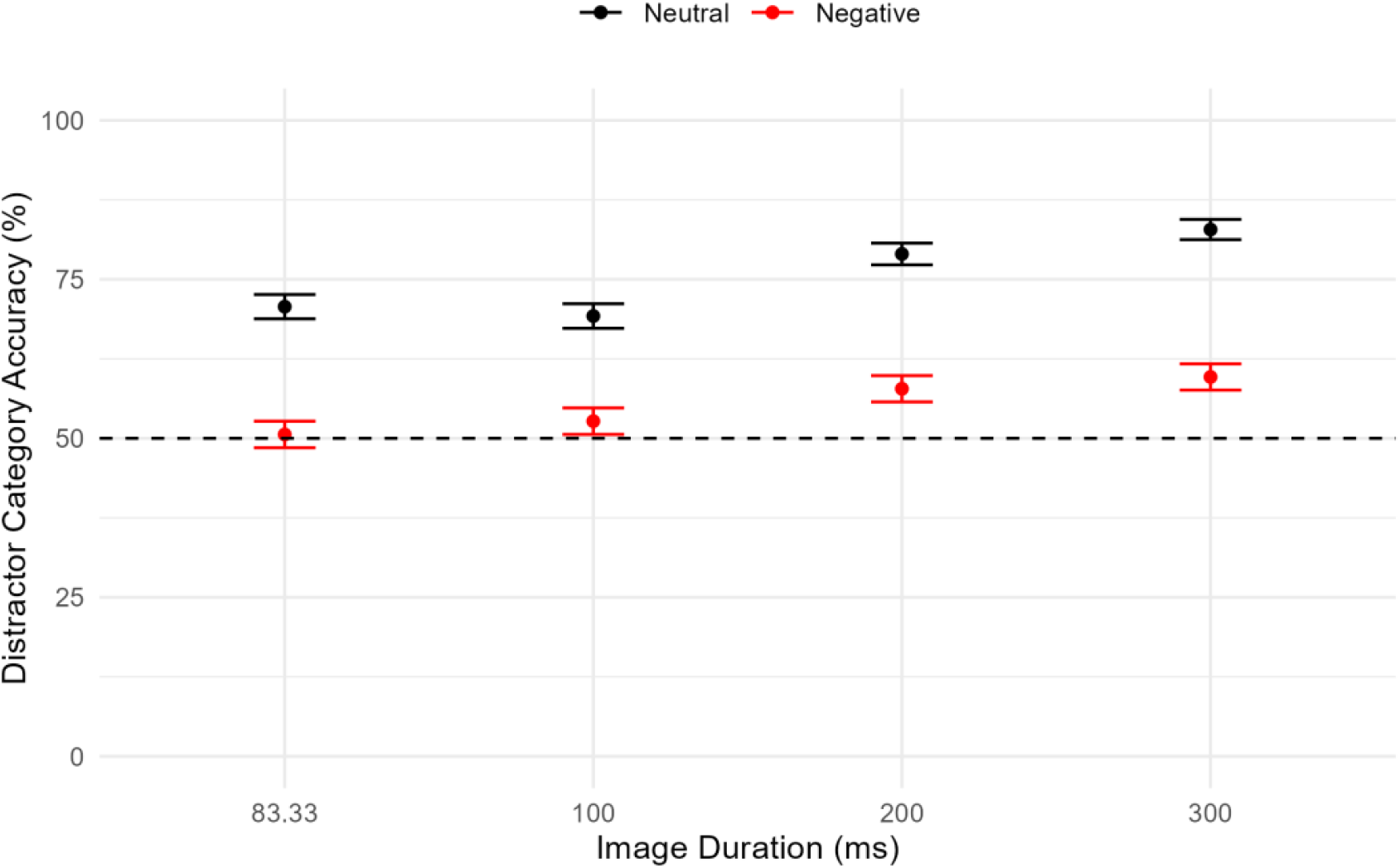
Accuracy of distractor category during recall phase by image duration and trial type (negative or neutral). Point and error bar represents mean ± SEM.

**Figure A6.**
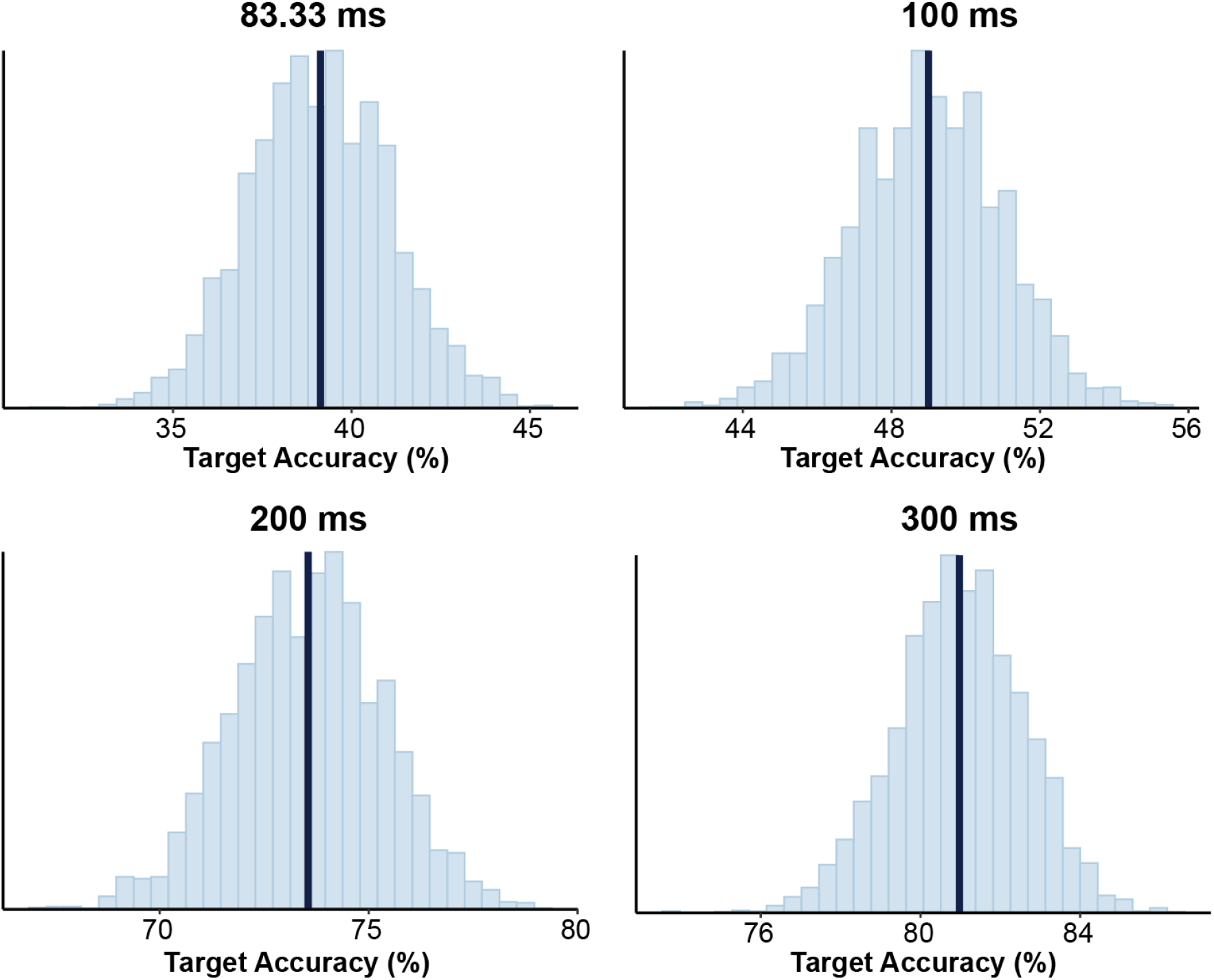
Experiment 3 observed average (black vertical lines) and simulated distribution (histogram) target accuracies for each image duration. Simulated data were created using model weights for estimating effect of negative distractors on target accuracy at each image duration condition.

**Figure A7.**
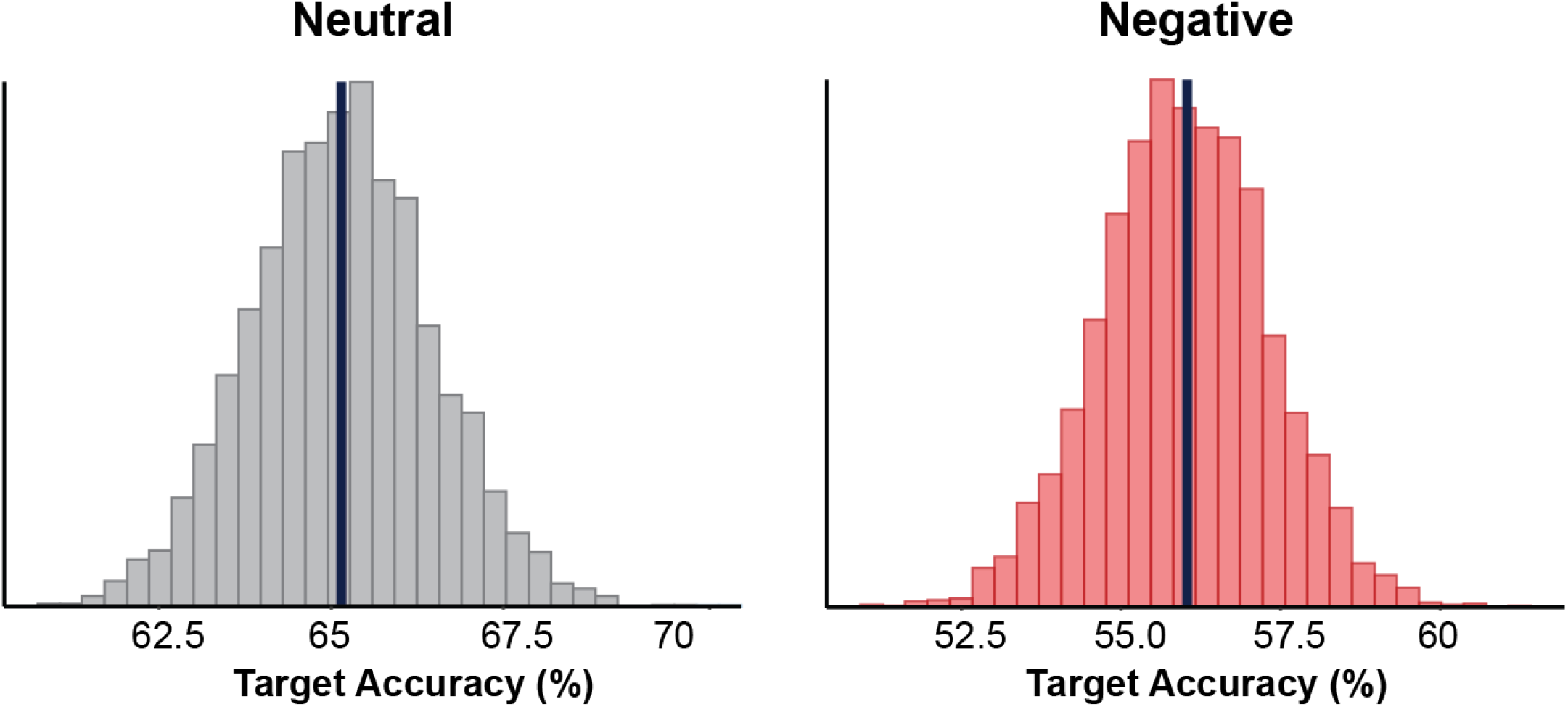
Experiment 3 observed average (black vertical lines) and simulated distribution (histogram) target accuracies for neutral (left) and negative (right) conditions.

